# A novel amidase signature family amidase from the marine actinomycete *Salinispora arenicola* CNS-205

**DOI:** 10.1101/342527

**Authors:** Ma Yanling, Zhang Xinfeng, Zeng Rong

## Abstract

We cloned a new gene from the amidase signature (AS) family, designated *am*, from the marine actinomycete *Salinispora arenicola* CNS-205. As indicated by bioinformatics analysis and site-directed mutagenesis, the AM protein belonged to the AS family. AM was expressed, purified, and characterised in *Escherichia coli* BL21 (DE3), and the AM molecular mass was determined to be 51 kDa. The optimal temperature and pH were 40 °C and pH 8.0, respectively. AM exhibited a wide substrate spectrum and showed amidase, aryl acylamidase, and acyl transferase activities. AM had high activity towards aromatic and aliphatic amides. The AM substrate specificity for anilides was very narrow; only propanil could be used as an effective substrate. The extensive substrate range of AM indicates it may have broad potential applications in biosynthetic processes and biodegradation.

## Introduction

Carboxylic acid amides can be hydrolysed by amidases (EC 3.5.1.4), forming carboxylic acids and ammonia. Most amidases also produce hydroxamic acids through their acyltransferase activity (Asano et al. 1982; Fournand et al. 1998). Amidases are very important for chemical industrial synthesis and for control of environmental pollution.

Amidases can be divided into two categories (Chebrou et al. 1996; Fournand and Arnaud 2001). The first category is the nitrilase superfamily, which is characterised by a cysteine residue and includes aliphatic amidases. The second category is the amidase signature (AS) family, which has a conserved GGSS signature in the amino acid sequence (Mayaux et al. 1990; Chebrou et al. 1996). Amidases are extensively present in bacteria, archaea and eukaryotes (d’Abusco et al. 2001; Galadari et al. 2006; Neu et al. 2007; Ohtaki et al. 2010; Politi et al. 2009).

The marine actinomycete *Salinispora arenicola* CNS-205 produces many bioactive natural products, including saliniketals A and B, which were originally isolated by Fenical and co-workers in 2006 (Fenical et al. 2006). Genome sequencing of the strain *S. arenicola* CNS-205 identified a gene encoding a putative amidase, named AM, which belongs to the AS family. The amidase activity of AM was confirmed, and its catalytic parameters and optimal conditions were determined. This enzyme was shown to have an abnormally wide substrate spectrum and activities.

## Materials and methods

### Chemicals

The chemicals used in this paper were graded as analytical reagents and purchased from J&K Scientific Company (Beijing, China).

### Bacterial strains and culture conditions

*S. arenicola* CNS-205 was cultured in liquid ISP2S (0.4% yeast extract, 1% malt extract, 0.4% glucose, and 7% sea salt; pH 7.3) in an incubator with rotation at 28 °C and 220 rpm and harvested after 2–3 days to obtain the genomic DNA.

### Cloning of the am gene

The genomic DNA of *S. arenicola* CNS-205 was extracted. Genes encoding potential AS family amidases were identified through BLASTP analysis (http://www.ncbi.nlm.nih.gov/blast). The primers used for the polymerase chain reaction of the *am* ORF were *am*-F (5′- GGGCATATGGCGGTGCAGGACATCA-3′) and *am*-R (5′- CAGGAATTCCAGTTTCGTCATGCCC-3′). The *Nde*I and *Eco*RI sites (underlined) were used to clone *am* into the protein expression vector pET-28a(+) (Novagen).

### AM expression and purification

*Escherichia coli* BL21 (DE3) carrying pET28a(+)-*am* was grown in Luria-Bertani medium with 100 μg ml^−1^ ampicillin at 37 °C (1 l medium was inoculated with 1% inoculum from a 20 ml overnight culture). The cultures were induced with 0.1 mM isopropyl-β-D-thiogalactopyranoside at OD_600_=0.6 and incubated for 20 h at 16 °C. Then, the cultures were centrifuged at 5,000 rpm for 10 min at 4 °C. The cell pellet was resuspended in prechilled binding buffer (20 mM Tris-HCl, pH 8.0, 0.5 mM NaCl, and 5 mM imidazole) and lysed by sonication on ice (60% amplitude, 4 s on and 10 s off). The supernatant was harvested after centrifugation at 15,000 rpm for 30 min at 4 °C. Then, the supernatant was loaded onto a His-Bind Ni resin column pre-equilibrated with binding buffer (GE Healthcare). An imidazole step gradient was used, and the His_6_-AM protein (50, 100, 200, 400, and 800 mM) was recovered in elution buffer (20 mM Tris-HCl, pH 8.0, and 0.5 mM NaCl). Then, SDS-PAGE and HPLC-MS (Thermo) were used to analyse the fractions with His_6_-AM. The supernatant containing His_6_-AM was further purified with PD-10 desalting columns (GE Healthcare) after it was concentrated through a VIVASPIN concentrator. The protein concentration was quantified with a NANODROP 2000c (Thermo), and a Thermo Hypersil GOLD C4 column (1.9 μ, 100×2.1 mm) was used. The recombinant proteins were assessed with HPLC-ESI-HRMS and eluted with a gradient of 0.1% formic acid (A) and CH3CN-containing 0.1% formic acid (B). The elution program was 2% B for 3 min, 2 to 20% B for 1 min, 20 to 70% B for 16 min, 70 to 90% B for 1 min, 90% B for 4 min, 90 to 2% B for 1 min, and 2% B for 4 min at a flow rate of 0.2 ml min^−1^. A Orbitrap mass spectrometer (Thermo) was used in positive ion mode, with scanning from m/z 300 to 2,000. Xcalibur software (v.1.1; Thermo Finnigan) was used for analysis, and the data were processed and deconvoluted.

### Amidase assays

The purified AM was resuspended in buffer (pH 8.0, 20 mM Tris-HCl, 10% glycerol and 100 mM NaCl). The assays were performed with 1 μg purified AM in 20 mM Tris-HCl, pH 8.0, 100 mM NaCl at 35 °C. The phenol-hypochlorite ammonia method (Weatherburn 1967) was used to assess the amidase activity, which yielded ammonia. The amount of enzyme catalysing the release of 1 μmol NH3/min was defined as one unit of enzyme activity. The Hanes-Woolf method was used to estimate the *K*_m_ and V_max_, and the *k*_cat_ and *k*_cat/_*K*_m_ values were determined, indicating a molecular mass of 51 kDa. Control reactions were performed without AM.

### Aryl acylamidase activity assay

The aryl anilide pesticides propanil, butachlor and acetochlor were assessed as substrates to determine the aryl acylamidase activity of AM. The aryl acylamidase activity was verified following the method of Shen et al. (2012) as follows: 1 μg of His_6_-AM was added to 0.2 mM anilide in 1 ml of 20 mM Tris-HCl, pH 8.0, 100 mM NaCl and incubated at 35 °C. Addition of HCl changed the pH to 3, and the sample was extracted with ethyl acetate, terminating the reaction. This organic layer was dried and re-dissolved in methanol. Reverse-phase HPLC (Shimadzu LC-20 AD, Waters 2998 photodiode array detector) with a Thermo C18 cartridge (particle size 3 μ; 2.1×150 mm) and 250 nm detection wavelength was used to recognise the reaction products, with 2:3 0.1% formic acid/methanol (isocratic elution mode) for 20 min at a flow rate of 0.2 ml min^−1^.

### Hydroxylamine-acyl transferase activity assay

The acyl transfer activity was detected as described by Fournand et al. (1998). All experiments were performed at 35 °C for 10 min, and the reaction system was as follows: 1 μg of AM, hydroxylamine hydrochloride (100 mM, pH 7.0), and amide or anilide (1∼25 mM) in buffer (20 mM Tris-HCl, pH 8.0, 100 mM NaCl). An acidic solution of FeCl3 (0.133 M in 0.68 M HCl) was adopted to terminate the reactions. The supernatant was initially centrifuged at 12,000 rpm for 10 min and subsequently collected, and the hydroxamate concentration was measured at λ=500 nm. The blank control experiments were performed without AM. The optical density of the experimental groups (marked A1) was determined. Additionally, the control group (marked A2) was assessed to determine the concentration (C) of hydroxamate C=(A1−A2)/εL (A refers to the optical density, ε denotes the coefficient of molar extinction, and L indicates the layer thickness). Different substrates have different ε values: propionamide, 1,029 M^−1^ cm^−1^; hydroxamate derivative of acetamide, 996 M−1 cm^−1^; propanil, 1,029 M^−1^ cm^−1^; isobutyramide, 1,016 M^−1^ cm^−1^. The control groups did not have AM. One unit of enzyme activity was defined as the amount of enzyme required to catalyse the formation or hydrolysis of 1 μmol of substrate or product every minute.

### Effects of temperature and pH on enzyme activity

For determination of the optimal temperature, the experiments were performed in buffer (20 mM Tris-HCl, pH 8.0, 100 mM NaCl) with 20 mM of the substrate benzamide. Thermo stability was detected by pre-incubation of the protein for 1 h at different temperatures. Then, the residual activity was tested at 35 °C. For determination of the optimal pH, the experiments were performed in various buffers as follows: 0.1 M sodium acetate buffer (pH 4.0, 4.5, 5.0, 5.5, and 6.0), 0.1 M potassium-phosphate buffer (pH 6.0, 6.5, 7.0, 7.5, 7.8, and 8.5), and 0.1 M Tris-HCl buffer (pH 7.5, 8.0, 8.5, 9.0, 9.5, and 10.0). For the pH stability detection, the experiments were performed in buffers with pH values that ranged from 4.0 to 10.0 with incubation at 25 °C for 1 h; then, the residual protein activity was tested at 35 °C.

### Impact of metal ions and other reagents

The impact of metal ions (Ni^2+^, Ba^2+^, Zn^2+^ and Ca^2+^) and chemical agents (1,10-phenanthroline, EDTA, SDS and PMSF) on the amidase activity was detected. The samples were preincubated for 10 min at 35 °C with benzamide as a substrate, and then, the amidase activity was determined as described previously.

### Site-directed mutagenesis of *am*

Site-directed mutagenesis primer pairs (Table 1) were designed to produce mutated *am* with a QuikChange site-directed mutagenesis kit (Stratagene). The recombinant plasmid pET-28a(+)-*am* served as a template in the mutagenesis reactions. The PCR products were purified by agarose gel electrophoresis, and the DNA bands with the appropriate sizes were eluted from the gel pieces. Then, the plasmid DNA was digested with *Dpn*I and transformed into competent *E. coli* BL21 (DE3) cells. Three mutant (K84A, S158A, and S182A) plasmids were constructed with this technique and were verified through DNA sequencing. The mutant proteins were expressed, purified, and analysed as described above.

**Table 1.**
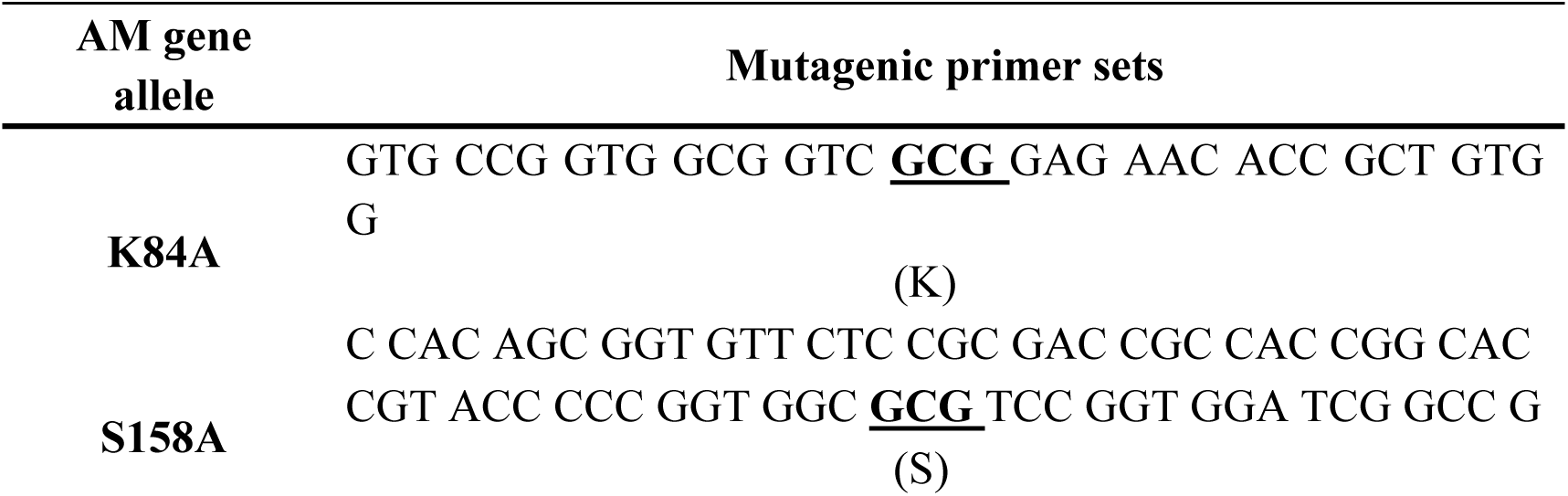
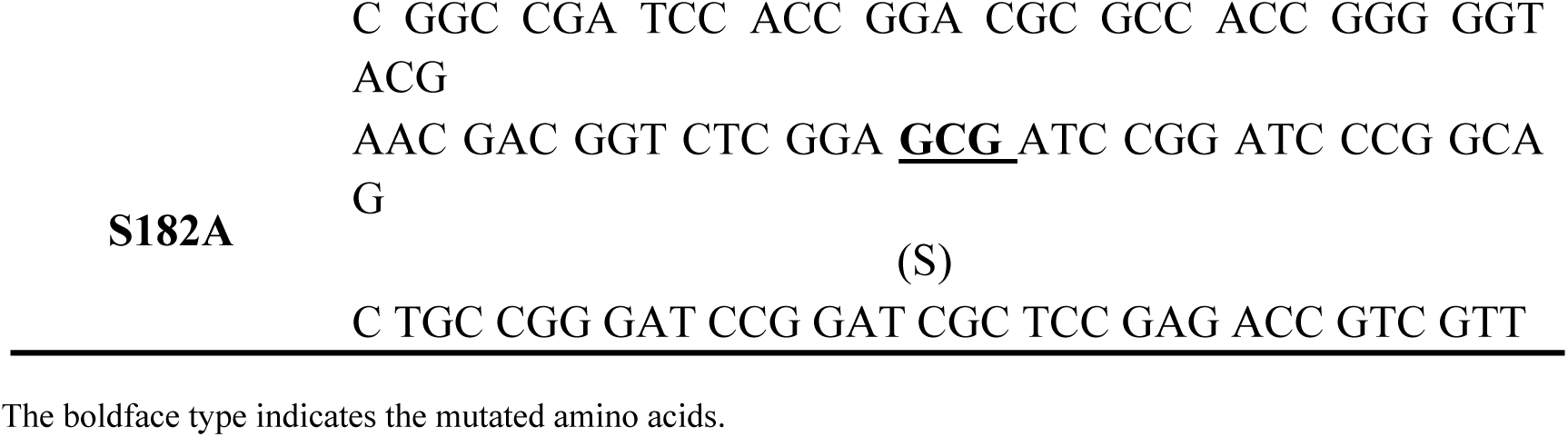
Oligonucleotides used for site-directed mutagenesis

### Nucleotide sequence accession number

The nucleotide sequence of *am* was stored in GenBank with a sequence ID of CP000850.1.

## Results

### Sequence analysis of the *am* gene

The amino acid sequence of AM was compared to sequences of known amidases accessible in the GenBank database. The comparisons showed that AM shared 29–45% identity with several enzymes, including an amidase from *Rhodococcus* sp. N771 (45% identity), an amidase from *Thermus thermophilus* HB8 (31% identity), the aspartyl/glutamyl-tRNA amidotransferase subunit A from *T. thermophilus* HB8 (33% identity), and the aspartyl/glutamyl-tRNA amidotransferase subunit A from *Thermotoga maritima* MSB8 (28% identity). Additionally, we confirmed the presence of Ser-Ser-Lys, the highly conserved catalytic triad of the AS family, in the enzyme amino acid sequence (Fig 1).

**Fig 1.**
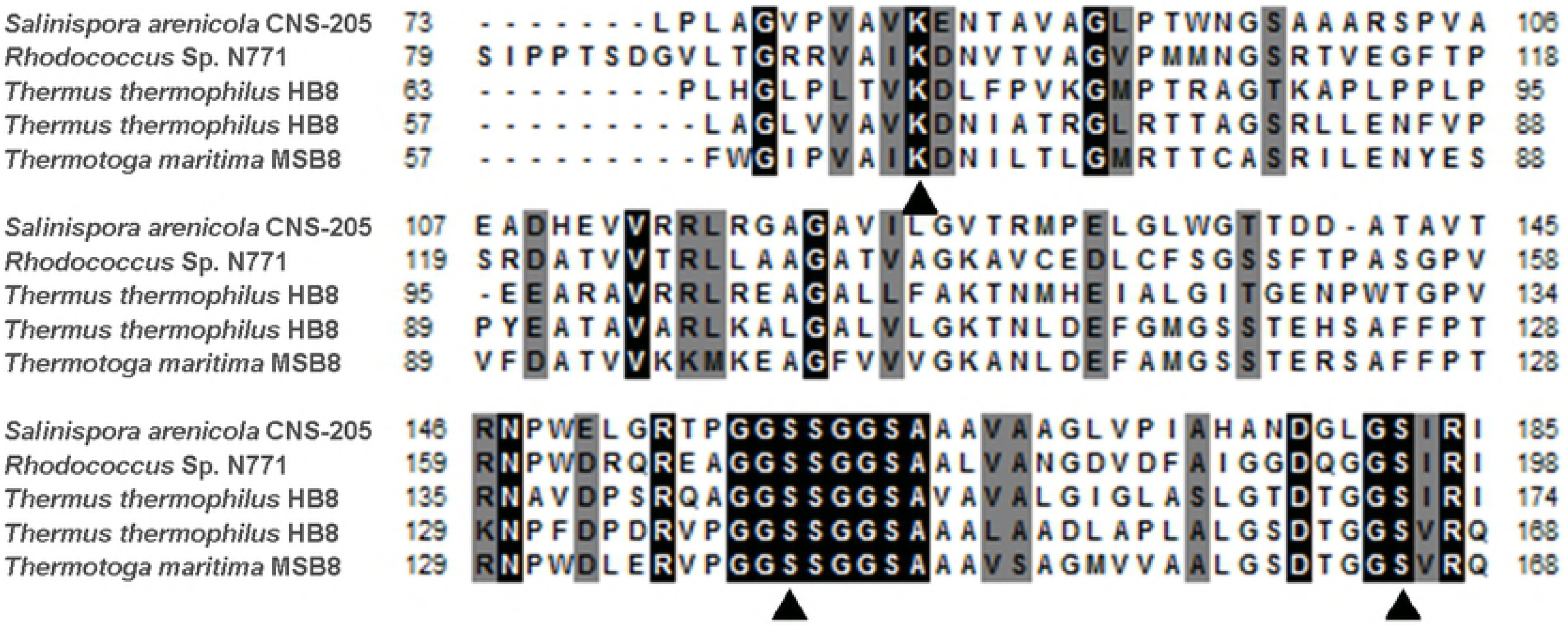
Comparison of the amino acid sequences of the AM and homologous proteins. Sequence alignment of the amino acid sequence of AM showing the high homology with the AS amidase (3A1K_A) from *Rhodococcus* Sp. N771, the *Thermus thermophilus* HB8 amidase (YP_145063.1), the *T. thermophilus* HB8 aspartyl/glutamyl-tRNA amidotransferase subunit A (YP_143839.1), and the *Thermotoga maritima* MSB8 aspartyl/glutamyl-tRNA amidotransferase subunit A (NP_229077.1). The alignment was produced by ClustalW. Multiple alignments were generated with BioEdit. The *dark grey* under the sequence indicates the residues of the Ser-Ser-Lys catalytic triad.

We demonstrated that this motif was the catalytic site through site-directed mutagenesis (Table 1). Thus, we showed that AM of *S. arenicola* CNS-205 contains the highly conserved catalytic triad Ser-Ser-Lys and is a member of the AS family.

### Expression and purification of AM

The fusion protein His_6_-AM was overexpressed in *E. coli* BL21 (DE3). The purity of the purified fusion protein was greater than 90%. The SDS-PAGE results indicated that the molecular mass of the major band was 51.2 kDa (Fig 2), which conformed to mass of the deduced protein sequence.

**Fig 2.**
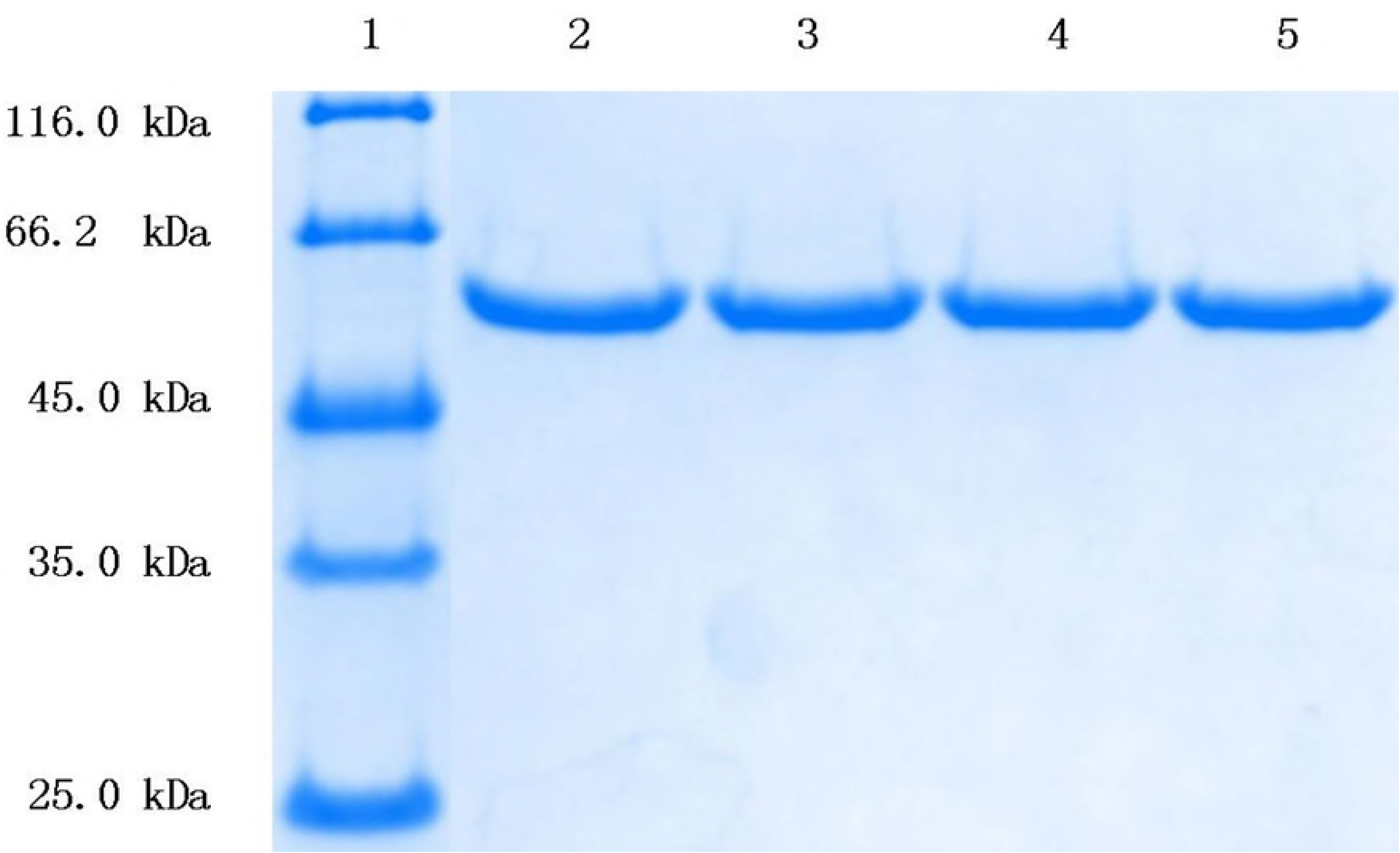
SDS-PAGE of AM-1 and its mutants. Lanes 1, protein molecular weight marker; 2, wild-type AM-1; 3, K84A; 4, S158A; and 5, S182A

To obtain the molecular mass of the proteins, we analysed the His_6_-AM fusion protein and three mutants (K84A, S158A, and S182A) by HPLC-ESI-HRMS. The molecular weights of the AM wild-type, K84A, S158A, and S182A were 51.037, 50.980, 51.021, and 51.021 kDa, which were consistent with the predicted values (Fig 3).

**Fig 3.**
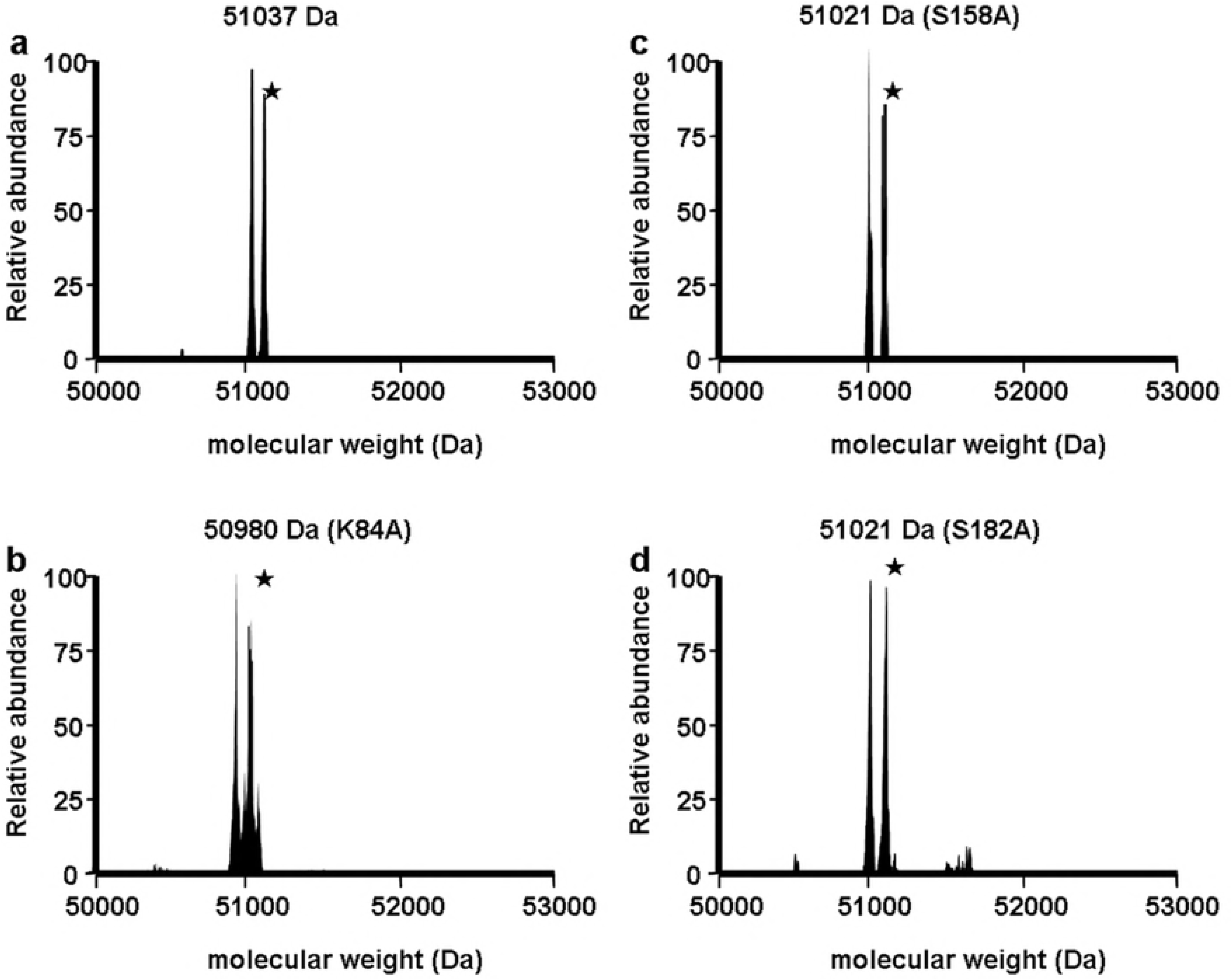
HPLC-ESI-HRMS analysis of AM. WT (a), K84A (b), S158A (c) and S182A (d). Extra minor peaks marked by asterisks denote glycosylation (+178 Da) of the N-terminal His_6_-tag added during the expression of the recombinant protein in *E*. *coli* (Geoghegan et al. 1999)

### Effects of temperature and pH on AM activity and stability

To determine the optimal temperature, we assessed the amidase activity at a temperature range from 15 to 65 °C with benzamide as the substrate. The AM activity peaked at 40 °C, and it showed an excessively wide peak (Fig 4a). More than 50% of the residual activity was observed at temperatures from 30 to 50 °C. Thermo stability tests indicated moderate loss of amidase activity within 1 h up to approximately 45 °C (Fig 4b). Only 9% of the activity was observed at 55 °C after 1 h, and the amidase activity was lost completely after 1 h at 60 °C.

**Fig 4.**
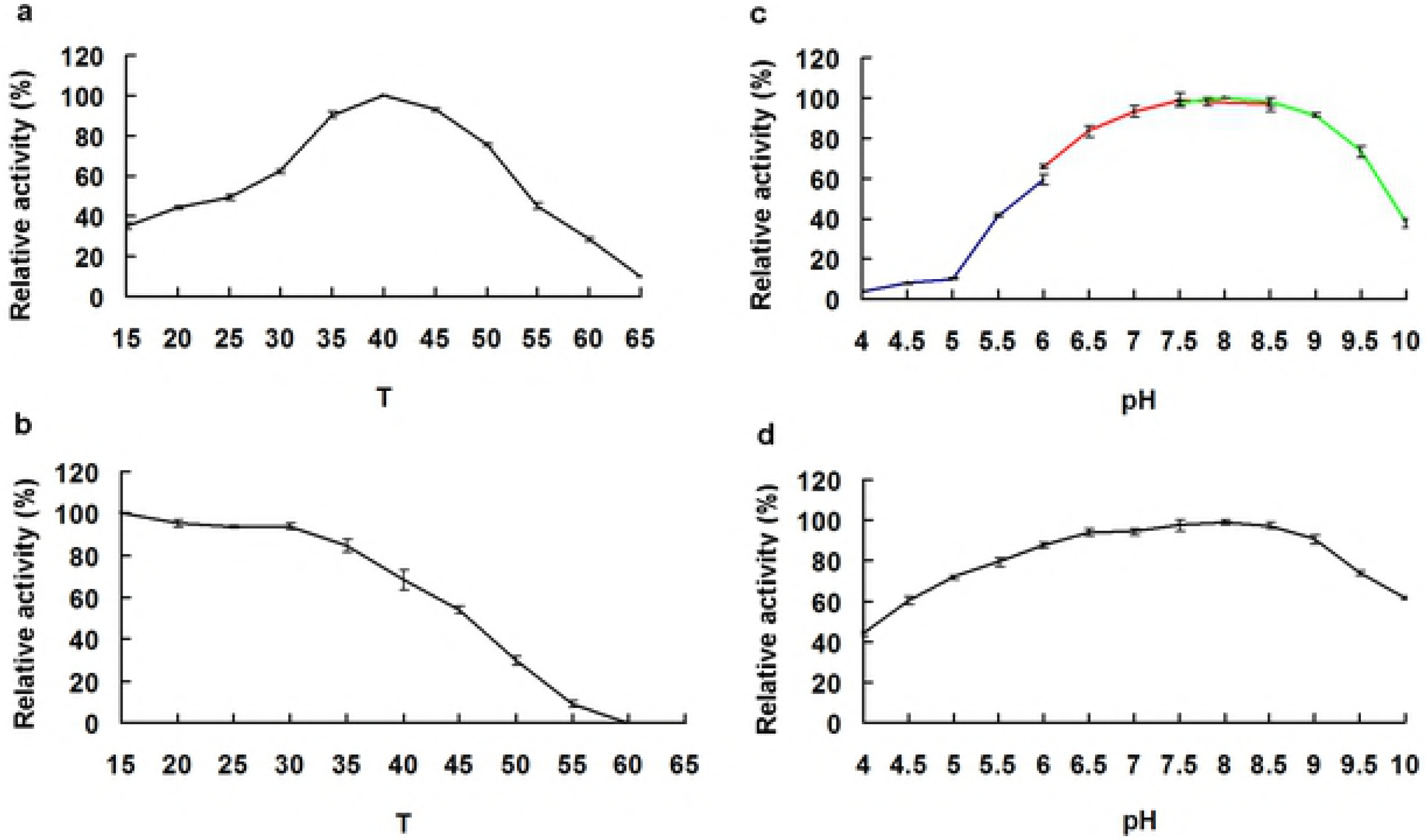
Effects of pH and temperature on AM activity and stability. **a** Determination of the optimal pH value. The reactions were performed at 35 °C for 10 min in buffers with varying pH values. **b** pH stability. The assays were performed in 20 mM Tris-HCl, 100 mM NaCl buffer (pH 8.0) at 35 °C for 10 min after pretreatment of the purified enzyme at 25 °C for 1 h in 0.1 M buffer (pH 4.0-10.0). **c** Determination of the optimal temperature. The activity was measured in 20 mM Tris-HCl, 100 mM NaCl, pH 8.0, at 15-65 °C for 10 min. **d** Thermal stability. The reactions were performed under optimal conditions after incubation of AM at the indicated temperature for 1 h.

The optimal pH for AM was determined in the buffers described in the Materials and methods. Figure 4c indicates that AM was highly active between pH 7.5 and 8.5. AM showed low activity below pH 4.5 or above pH 10.0. For the pH stability test, AM was preincubated for 1 h at different pH values, and the results indicated that more than 60% residual activity was observed between pH 4.5 and 10.0 (Fig 4d).

### Impact of metal ions and other reagents

The majority of the metal ions, including Ba^2+^, Ca^2+^, Zn^2+^, and Ni^2+^, in the assays exerted no noticeable effect on the amidase activity of AM (Table 2).

**Table 2.** Effects of metal ions and inhibitors on the amidase activity of AM

| Reagent<br>(1 mM) | Relative activity<br>(%) | Reagent<br>(1 mM) | Relative activity<br>(%) |
| --- | --- | --- | --- |
| No addition | 100 | PMSF | 10.13 ± 0.60 |
| Ba<sup>2+</sup> | 94.30 ± 2.59 | 1,10-Phenanthroline | 82.46 ± 0.73 |
| Ca<sup>2+</sup> | 88.60 ± 2.13 | EDTA | 77.64 ± 0.92 |
| Zn<sup>2+</sup> | 85.09 ± 3.87 | SDS | 51.78 ± 0.27 |
| Ni<sup>2+</sup> | 96.93 ± 1.90 |  |  |

The activity was strongly inactivated by PMSF, which is an inhibitor of serine hydrolases. However, the chelating agents EDTA and 1,10- phenanthroline (10 mM) resulted in only a 20–30% inhibition of AM hydrolysis, demonstrating that these chemicals did not chelate a possible divalent cation(s) required for the activity of the enzyme. The surfactant SDS showed a 48.22% inhibition of AM activity.

### Substrate spectrum

To determine the substrate specificity of AM, we assessed whether the purified AM could hydrolyse different aromatic and aliphatic amides. The results showed that AM had high activity towards the aromatic and aliphatic amides, including acetamide, propionamide, propanil, benzeneacetamide and benzamide (Table 3).

**Table 3.** Substrate spectrum of AM

| Substrate | Relative activity <sup>a</sup><br>(%) | Substrate | Relative activity <sup>a</sup><br>(%) |
| --- | --- | --- | --- |
| Acetamide | 103.79 ± 0.46 | Propanil | 95.91 ± 0.42 |
| Benzamide | 100 | Propanamide | 106.36 ± 0.38 |
| Benzeneacetamid | 107.75 ± 0.48 | Isobutyramide | 18.02 ± 0.72 |
| Nicotinamide | 56.42 ± 1.29 | Acetochlor | 0 |
| Pyrazinamide | 43.01 ± 2.76 | Butachlor | 0 |

<sup>a</sup> The activity with benzamide was defined as 100%. All measurements were performed in triplicate

The aromatic amides, including nicotinamide and pyrazinamide, with substitutions of one or two carbons in the ring by a nitrogen, had a negative impact on the activity. The anilide substrate range of AM was very narrow, and the protein could not hydrolyse butachlor and acetochlor. Only propanil was a good substrate for AM.

Kinetic parameters for AM were estimated by the Hanes-Woolf method. The *K*_m_ values for acetamide and propionamide were 3.36±0.17 mM and 3.33±0.08 mM (Table 4).

**Table 4.** Kinetic parameters for AM amidase reaction

| Substrate | $K_m$ (mM) | $V_{max}$<br>(mM min<sup>-1</sup>) | $k_{cat}$ (s<sup>-1</sup>) | $k_{cat}/K_m$<br>(mM<sup>-1</sup> s<sup>-1</sup>) |
| --- | --- | --- | --- | --- |
| Acetamide | $3.36 \pm 0.17$ | $0.491 \pm 0.001$ | $63.89 \pm 0.20$ | $19.06 \pm 0.94$ |
| Nicotinamide | $6.20 \pm 0.04$ | $0.293 \pm 0.006$ | $38.19 \pm 0.75$ | $6.16 \pm 0.12$ |
| Propionamide | $3.33 \pm 0.08$ | $0.503 \pm 0.001$ | $65.54 \pm 0.08$ | $19.69 \pm 0.43$ |
| Benzamide | $3.49 \pm 0.12$ | $0.508 \pm 0.005$ | $66.15 \pm 0.66$ | $18.95 \pm 0.72$ |

<sup>a</sup> All substrates were tested at 25-200 mM, except for benzamide, which was insoluble above 50 mM

The anilide substrate range of AM was very narrow, and the protein could not hydrolyse acetochlor and butachlor. Only propanil was a good substrate for AM, but the *K*_m_ value could not be determined due to the low solubility of the compound.

### Site-directed mutagenesis

The potential catalytic active site residues of AM were replaced by the QuikChange site-directed mutagenesis kit. The mutants were overexpressed in *E. coli* BL21 (DE3) cells and further purified as described above. The results indicated that the AM K84, S158 and S182 mutants had no activity with benzamide as a substrate. These results suggest that AM is a member of the AS family and utilises the highly conserved catalytic triad Ser-Ser-Lys.

## Discussion

An isolate of *S. arenicola* CNS-205 identified by Fenical (Fenical et al. 2006) and co-workers in 2006 encodes a putative amidase. Sequence alignments of the primary AM sequence indicated that AM had a high similarity with the AS family and showed that AM contains the central GGSS signature, which is a typical characteristic of the AS family. The point mutation results also indicated that no hydrolytic activity could be detected in the K84A, S158A, and S182A mutants. These findings indicated that AM belongs to the AS family.

The effects of different metal ions and chemical reagents on AM activity were different. AM activity was affected by reducing agents, such as PMSF, and the results revealed that serine was the active site of the amidase. This result was consistent with the crucial role of Ser^183^, as revealed by site-directed mutagenesis experiments. A metal chelating agent (EDTA) did not impact the activity, indicating that a possible divalent cation(s) required for enzyme activity was not chelated by these chemicals.

Analysis of the substrate specificity of AM showed that the enzyme had high activity against short-chain aliphatic amide substrates (acetamide, isobutyramide and propanamide), which are typical substrates of the AS family. Interestingly, AM also hydrolyses ring amide substrates, such as aromatic and heterocyclic amides. The hydrolytic product of nicotinamide is nicotinic acid, a water-soluble B-complex vitamin, which has been extensively applied in treatment of schizophrenia, autoimmune diseases, hypercholesterolemia, diabetes and osteoarthritis. Benzoic acid, the hydrolytic product of benzamide, has antifungal activity and is extensively used as a preservative in production of processed and convenience foods.

AM also had aryl acylamidase activity against aniline substrates, including propanil (a commercial amide-containing pesticide), which was hydrolysed to produce 3,4-dichloroaniline. However, acetochlor and butachlor, which are structurally analogous to propanil, were not substrates for AM, indicating the anilide substrate range of AM was very narrow. Propanil, an acyl anilide herbicide, can contaminate the soil environment, and AM, by attacking the amide bonds in propanil, can reduce its concentrations in soil. Thus, AM may have potential applications in bioremediation.

In addition to the amidase and aryl acylamidase activities, acyl transferase activity is an important characteristic as it produces hydroxamic acids (Fournand et al. 1998). In this study, AM from *S. arenicola* CNS-205 had acyl transferase activity on anilide substrates, including propanil. The extensive substrate specificity range and acyl transferase activity indicate that AM has broad potential applications in biosynthesis processes and biodegradation.

Overall, a new amidase gene, AM, was cloned from *S. arenicola* CNS-205, and the amidase, aryl acylamidase, and acyl transferase activities of the enzyme were verified. These activities indicate that AM has a broad substrate spectrum. AM may be a potential agent for environmental remediation and for the biosynthesis of novel amides by virtue of these characteristics, as well as the broad pH tolerance of the enzyme.

